# CCR2 Regulates Vaccine-Induced Mucosal T-Cell Memory to Influenza A Virus

**DOI:** 10.1101/2021.03.24.436901

**Authors:** Woojong Lee, Brock Kingstad-Bakke, Ross M. Kedl, Yoshihiro Kawaoka, M. Suresh

## Abstract

Elicitation of lung tissue-resident memory CD8 T cells (T_RM_s) is a goal of T-cell based vaccines against respiratory viral pathogens such as influenza A virus (IAV). Chemokine receptor 2 (CCR2)-dependent monocyte trafficking plays an essential role in the establishment of CD8 T_RM_s in lungs of IAV-infected mice. Here, we used a combination adjuvant-based subunit vaccine strategy that evokes multifaceted (T_C_1/T_C_17/T_H_1/T_H_17) IAV nucleoprotein-specific lung T_RM_s, to determine whether CCR2 and monocyte infiltration are essential for vaccine-induced T_RM_ development and protective immunity to IAV in lungs. Following intranasal vaccination, neutrophils, monocytes, conventional dendrtitic cells (DCs) and monocyte-derived DCs internalized and processed vaccine antigen in lungs. We also found that Basic Leucine Zipper ATF-Like Transcription Factor 3 (BATF-3)-dependent DCs were essential for eliciting T cell responses, but CCR2 deficiency enhanced the differentiation of CD127^HI^/KLRG-1^LO^, OX40^+ve^CD62L^+ve^ and mucosally imprinted CD69^+ve^CD103^+ve^ effector and memory CD8 T cells in lungs and airways of vaccinated mice. Mechanistically, increased development of lung T_RM_s, induced by CCR2 deficiency was linked to dampened expression of T-bet, but not altered TCF-1 levels or T cell receptor signaling in CD8 T cells. T1/T17 functional programming, parenchymal localization of CD8/CD4 effector and memory T cells, recall T cell responses and protective immunity to a lethal IAV infection were unaffected in CCR2-deficient mice. Taken together, we identified a negative regulatory role for CCR2 and monocyte trafficking in mucosal imprinting and differentiation of vaccine-induced T_RM_s. Mechanistic insights from this study may aid the development of T-cell-based vaccines against respiratory viral pathogens including IAV and SARS-CoV-2.

**Importance:** While antibody-based immunity to influenza A virus (IAV) is type and sub-type specific, lung and airway-resident memory T cells that recognize conserved epitopes in the internal viral proteins are known to provide heterosubtypic immunity. Hence, broadly protective IAV vaccines need to elicit robust T-cell memory in the respiratory tract. We have developed a combination adjuvant-based IAV nucleoprotein vaccine that elicits strong CD4 and CD8 T cell memory in lungs and protects against H1N1 and H5N1 strains of IAV. In this study, we examined the mechanisms that control vaccine-induced protective memory T cells in the respiratory tract. We found that trafficking of monocytes into lungs might limit the development of anti-viral lung-resident memory T cells, following intranasal vaccination. These findings suggested that strategies that limit monocyte infiltration can potentiate vaccine-induced frontline T-cell immunity to respiratory viruses such as IAV and SARS-CoV-2.

## Introduction

Upon respiratory infection, conventional dendritic cells (cDCs) endocytose and process antigens in the pulmonary environment and migrate to the draining lymph nodes (DLNs) to stimulate effector CD8 T cell responses (1–3). Naïve T cells recognize antigens in context of antigen-presenting cells (APCs), and undergo a distinct program of proliferation and differentiation into effector T cells in lung-draining lymph nodes, which traffic to lungs and clear the infection (4, 5). Upon trafficking to the lung tissues, effector T cells may encounter another round of antigenic stimulation by pulmonary APCs, including conventional dendritic cells (cDCs), monocyte-derived DCs, alveolar macrophages, monocytes, and neutrophils (6–9). In addition to antigenic re-stimulation, inflammatory milieu may dictate differentiation and functional diversification of effector CD8 T cells in the lung, which in turn might regulate the quality, quantity, anatomical localization and durability of T cell memory (10–12). Upon clearance of pathogens, phenotypically diverse memory CD4/CD8+ T cells persist in the lungs. Some effector T cells within the lung differentiate into a subset of tissue-resident memory T cells (T_RM_) that can permanently reside within the lungs or migrate into draining lymph nodes (13); lung- and airway-resident CD8 T_RM_s are crucial for providing broad heterosubtypic immunity against influenza (14–17). Additionally, some effector T cells differentiate into memory T cells that circulate between lymph and blood (T_CM_) or between blood and peripheral tissues (T_EM_) (18–20).

To protect individuals from respiratory pathogens such as influenza A viruses (IAV) and SARS-CoV-2, vaccines need to engender balanced humoral and T cell-mediated immunity (21–26). While establishment of lung T_RM_s is likely one of the major goals of designing effective T cell-based vaccines against respiratory pathogens, it has been challenging to elicit durable and effective mucosal T-cell immunity in the lungs using currently available vaccine platforms (27, 28). While many of the current FDA-approved vaccines are administrated by intramuscular or subcutaneous injections, there is emerging interest in designing intranasal vaccines, which can be directly delivered to the mucosal surface of respiratory tract. Intranasally administered vaccines could be more effective than injected vaccines, because intranasal vaccination can evoke virus-specific antibodies and memory CD8 T cells in the upper respiratory tract that can expeditiously clear the pathogens at the portal of entry. Hence, identifying safe and effective mucosal adjuvants is likely crucial to mitigate the global impact of currently circulating and newly emerging respiratory pathogens, such as SARS-CoV-2.

Understanding key cellular interactions that regulate the generation and persistence of memory T cell subsets is vital for designing effective vaccines. Several factors govern the development of CD8 T_RMs_ following an infection and the roles of regulatory cytokines (e.g. IL-15, and TGF-β) and antigenic stimulation have been extensively investigated in recent years (10, 29–33). There is good evidence that pulmonary monocytes interact with effector CD8 T cells in the lung to drive T_RM_ differentiation following vaccinia or influenza infection (34, 35), but the underlying mechanisms are unknown. Further, it is unclear whether cellular and molecular factors that regulate T_RM_ formation during a viral infection play similar roles in the development of T_RM_s following vaccination. Given the importance of T_RM_ for protective immunity against respiratory viruses, it is important to elucidate whether monocytes play an important role in engendering T cell immunity following vaccination. Insights from such studies might aid in the rational development of adjuvants that can drive potent vaccine-induced T cell responses by engaging monocytes in the lungs.

Adjuplex (ADJ) is a carbomer-based nano-emulsion adjuvant that is known to elicit robust neutralizing antibodies to malarial and HIV envelope glycoproteins in mice and non-human primates (36, 37). We have previously reported that subunit protein formulated in ADJ protects against vaccinia virus and IAV in mice by enhancing DC cross-presentation (38, 39). Additionally, we demonstrated that ADJ, in combination with Toll-like receptor 4 (TLR4) agonist glucopyranosyl lipid A (GLA), induces robust effector and T_RM_ CD8 and CD4 T cell responses to IAV nucleoprotein antigen and engenders effective T cell-dependent protection against H1N1 and H5N1 IAVs (40). However, mechanisms underlying the development of vaccine-induced protective CD4/CD8 T_RM_s in the respiratory tract, remain largely unknown. In this study, in mice administered with a subunit vaccine formulated in ADJ+GLA, we have examined the identity and kinetics of the antigen-processing cell types in lungs and DLNs, and then assessed the role of CCR2 and monocytes in orchestrating the differentiation of effector and memory CD8/CD4 T cells. Further, we examined whether programming of recall T cell responses and protective immunity are affected by CCR2 deficiency. We found that CCR2 play key roles in promoting terminal differentiation of effector T cells and limiting CD103, OX40 and CD62L expression on effector and memory CD8 T cells. Despite altered differentiation of effector and memory T cells, programming of recall T cell responses and T cell-dependent protective immunity were unaffected in CCR2-deficient mice. These findings provided unique insights into immunological mechanisms that orchestrate memory T cell differentiation following mucosal vaccination against respiratory viral infection.

## RESULTS

### Dynamics of antigen-processing innate immune cells in lungs and draining lymph nodes and the role of BATF3-dependent DCs in T cell responses to intranasal vaccination with a subunit protein antigen

To reiterate, intranasal (IN) vaccination with a subunit protein formulated with a combination adjuvant (Adjuplex [ADJ] + TLR4 agonist glucopyranosyl lipid A [GLA]) elicited high numbers of T_RM_ CD8 T cells and provided robust protection against IAV (40). To better understand the role and identity of antigen-processing innate immune cells in eliciting a strong T_RM_ response, we vaccinated mice IN with DQ-OVA formulated in ADJ+GLA; only upon proteolytic digestion, DQ-OVA emits green (DQ green) or red fluorescence (DQ red) **(Figure 1A)**. At days 2, 5 and 8 after vaccination, we quantified DQ green^+^/red^+^ innate immune cell subsets that contained processed DQ-OVA in lungs and DLNs **(Figure 1B and Supplementary Figure 1A).** The percentages of DQ green^+^/red^+^ cells were highest at day 2, but dwindled by day 8 after vaccination. At days 2 and 5 after vaccination, neutrophils, monocytes and monocyte-derived DCs constituted a major proportion of cells containing processed OVA in lungs (**Figure 1C**). Notably, between days 2 and 8 after vaccination, the percentages of processed DQ-OVA-bearing CD103^+ve^ migratory DCs increased both in the lungs and draining lymph nodes. By day 8, DQ-OVA was predominantly detected in CD103^+ve^ DCs in DLNs (**Figure 1D**).

**Figure 1.**
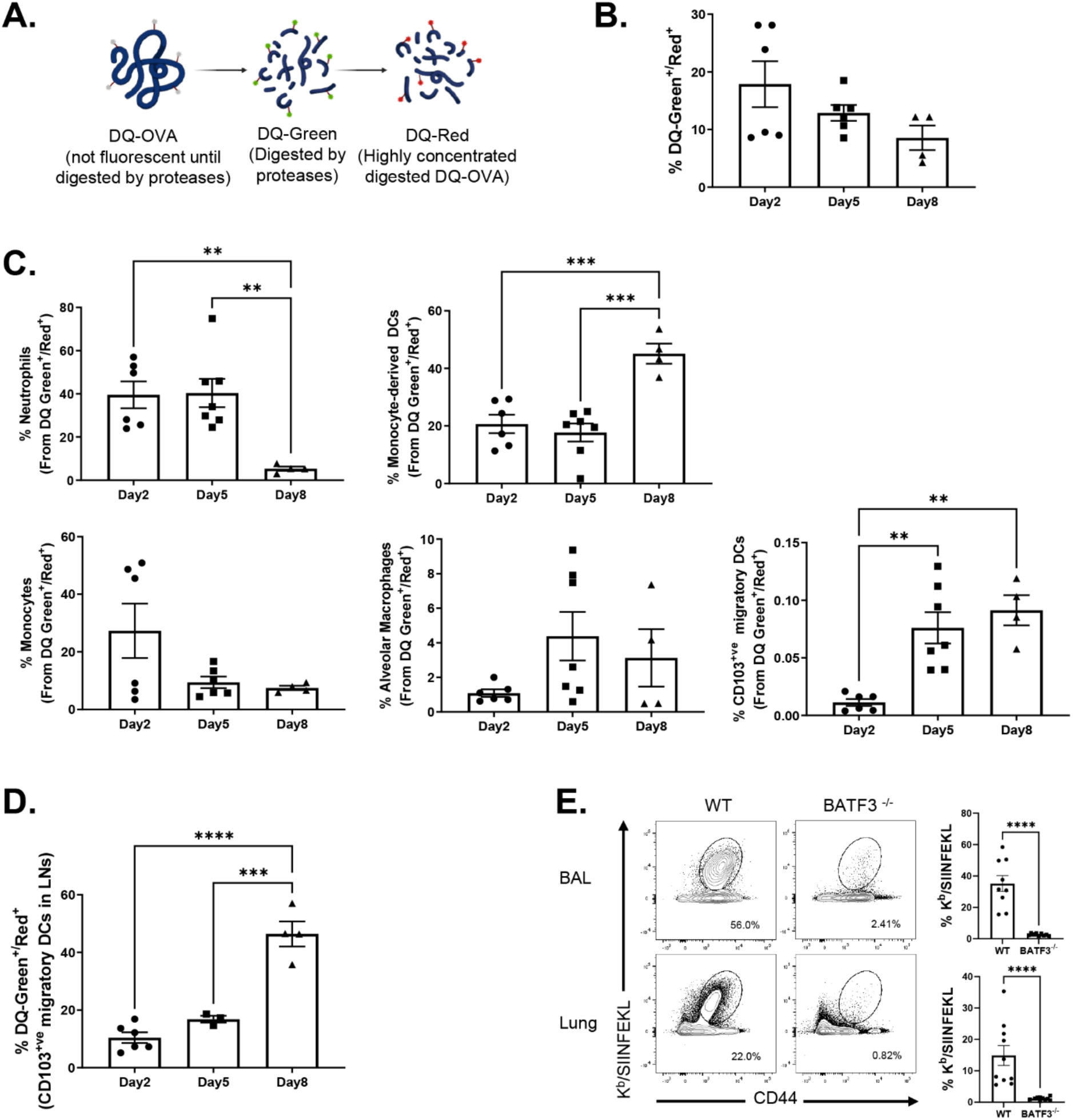
Dynamics of antigen processing by innate immune cells following intranasal vaccination. (A) Cartoon shows mechanism of action of DQ-OVA. (B) Groups of C57BL/6 mice were vaccinated IN with DQ-OVA protein formulated in ADJ (5%) and GLA (5ug). At day 2, 5 and 8 after immunization, single-cell suspensions of lung cells were stained with anti-CD11b, anti-Siglec-F, anti-CD11c, anti-CD64, anti-Ly6G, anti-Ly6C, anti-CD103, and anti I-A/I-E and immunophenotyped. (B) Single cell suspension of lungs were analyzed for processed DQ-OVA (green and red fluorescence [DQ-Green^+ve^/Red^+ve^]). (C) Percentages of innate immune cell subsets amongst total DQ-Green^+ve^/Red^+ve^ cells. (D) Percentages of DQ-green^+ve^ and DQ-red^+ve^ among CD103+ migratory DCs in LNs. Data are pooled from two independent experiments. (E) Wild-type (WT) and BATF3-deficient (BATF3^-/-^) mice were vaccinated intranasally with OVA (10ug) formulated in ADJ (5%) + GLA (5ug). On the 8th day after vaccination, the total number of activated OVA SIINFEKL-specific CD8 T cells in the lung and BAL were quantified by staining lung cells with K^b^/SIINFEKL tetramers, anti-CD8 and anti-CD44; FACS plots are gated on total CD8 T cells, and the numbers are percentages of tetramer-binding cells among gated CD8 T cells. The data represent one of two independent experiments (B-H). Mann-Whitney U test, *, **, and *** indicate significance at *P*<0.1, 0.01 and 0.001 respectively

Previous work has shown that development of migratory CD103^+ve^ DCs is dependent upon the transcription factor BATF3, and T cell responses elicited by cross-presenting DCs are compromised in BATF3-deficient (BATF3^-/-^) mice (41, 42). To assess whether BATF3-dependent migratory DCs are required to elicit CD8 T cell responses, we vaccinated wild type (WT) and BATF3^-/-^ mice with OVA formulated in ADJ+GLA. At day 8 after vaccination, we quantified OVA SIINFEKL epitope-specific CD8 T cells in lungs using MHC I tetramers (**Figure 1E**). High numbers of SIINFEKL-specific CD8 T cells accumulated in lungs of WT mice, but the numbers of such cells were substantively reduced in lungs of BATF3^-/-^ mice. These findings suggested that elicitation of CD8 T cell response by ADJ+GLA requires BATF3 and likely BATF3-dependent migratory DCs.

### Role of pulmonary monocytes in mucosal imprinting and differentiation of vaccine-induced effector CD8 T cells in the respiratory tract

Studies of T cell responses to IAV in CCR2-deficient (CCR2-/-) mice have suggested that recruitment of monocytes into lungs might play a key role in development and maintenance of T_RM_s in the respiratory tract (34, 35). Data in **Figure 1 A** showed that high percentages monocytes and monocyte-derived DCs internalized and processed protein antigen in lungs, following IN vaccination. Therefore, it was of interest to determine whether monocytes and monocyte-derived DCs regulated CD8 T cell responses to ADJ+GLA-adjuvanted subunit vaccine. We immunized WT and CCR2^-/-^ mice IN twice (at 3-week interval) with influenza virus nucleoprotein (NP) formulated with ADJ/GLA. At day 8 after booster vaccination, we quantified the percentages and number of NP366-specific CD8 T cells in airways (bronco-alveolar lavage [BAL]) and lungs. Despite the absence of monocytes and monocyte-derived DCs (not shown), the percentages and numbers of NP366-specific CD8 T cells in lungs and airways of CCR2-/- mice were comparable to those in WT mice **(Figure 2).** These data suggested that CCR2 and pulmonary monocytes are not essential for CD8 T cell responses to vaccination with ADJ+GLA.

**Figure 2.**
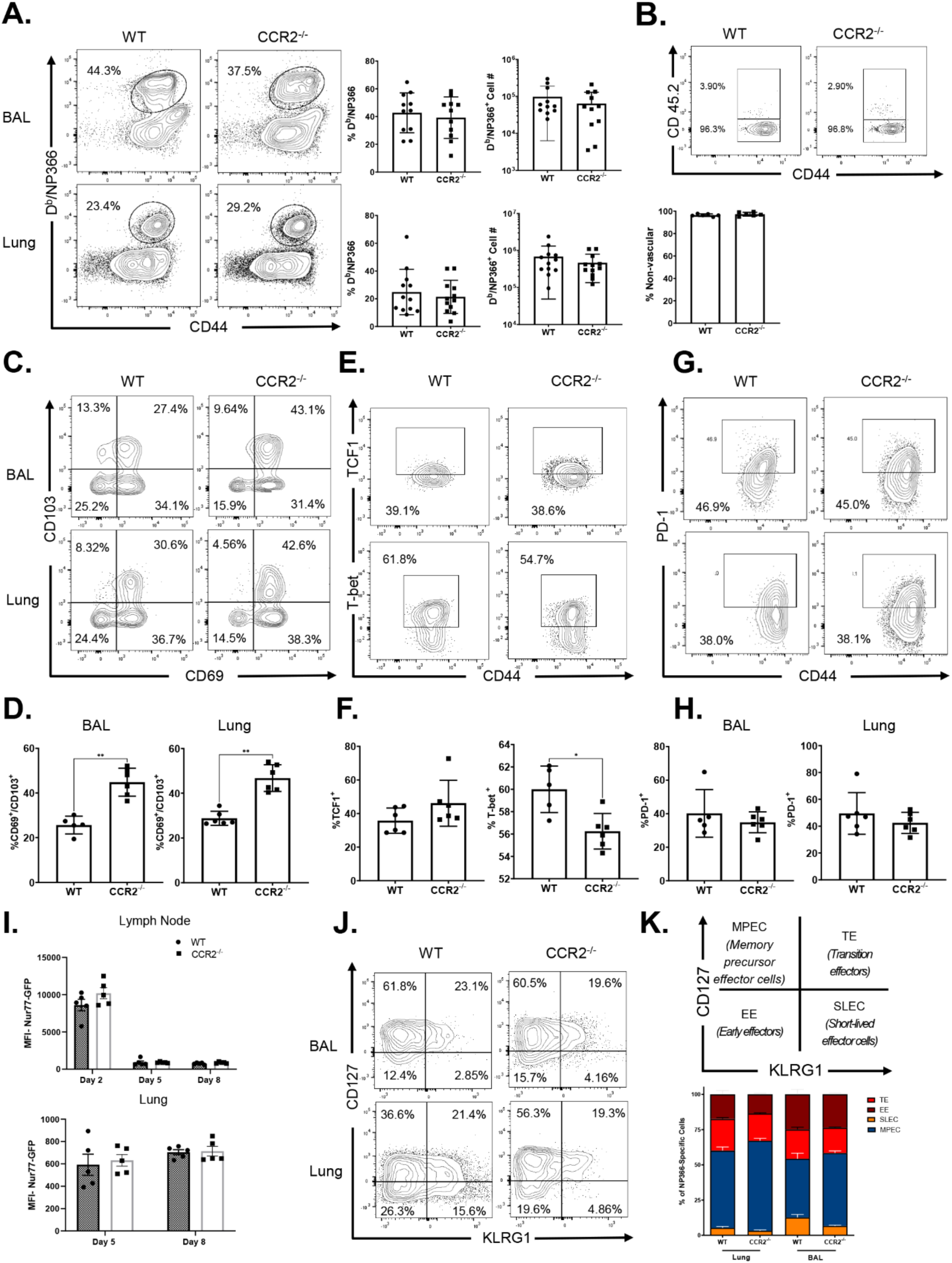
Effector CD8 T cell response to adjuvanted subunit vaccine in CCR2^-/-^ mice. Wild Type (WT) or CCR2^-/-^mice were immunized intranasally (IN) twice (21 day apart) with Influenza A H1N1 Nucleoprotein (NP) formulated in ADJ (5%) and GLA (5ug). To distinguish non-vasicular cells from vasicular cells in the lungs, mice were injected intravenously with fluorescent-labeled anti-CD45.2 antibodies, 3 min prior to euthanasia (CD45.2^+ve^ – vascular; CD45.2^-ve^ – non-vascular.) At day 8 post-vaccination, single-cell suspensions prepared from the lungs and bronchoalveolar lavage (BAL) were stained with viability dye, followed by D^b^/NP366 tetramers in combination with anti-CD4, anti-CD8, amti-CD44, anti-CD69, anti-CD103, anti-PD-1, anti-KLRG1, anti-CD127, anti-T-bet and anti-TCF-1. (A) FACS plots show percentages of tetramer-binding cells among CD8 T cells. (B) Percentages of vascular and non-vascular cells in NP366-specific CD8 T cells. FACS plots are gated on tetramer-binding CD8 T cells. (C, E, G) FACS plots are gated on D^b^/NP366 tetramer-binding CD8 T cells and numbers are percentages of CD69^+^CD103^+^, PD-1^+^, T-bet^+^ and TCF-1^+^ cells in respective gates or quadrants. (I) Naïve CD45.1^+^ Nur77-eGFP OT-I CD8 T cells were adoptively transferred into congenic CD45.2 WT or CCR2^-/-^ mice. Twenty four hours after cell transfer, mice were vaccinated IN with OVA formulated with ADJ (5%) and GLA (5ug). At days 2, 5, or 8 post-vaccination, single-cell suspensions from mediastinal lymph nodes and lungs were stained with anti-CD8, anti-CD45.1 antibodies, and K^b^/SIINFEKL tetramers. The MFIs of Nur77-eGFP in donor CD45.1^+ve^ OT-I CD8 T cells were quantified by flow cytometry. (J) FACS plots are gated on tetramer-binding CD8 T cells and numbers are percentages of effector subsets shown in K. Data are pooled from two independent experiments (A) or represent one of two independent experiments (B-K). Mann-Whitney U test, *, **, and *** indicate significance at P<0.1, 0.01 and 0.001 respectively.

To determine whether CCR2 deficiency altered the localization of vaccine-elicited NP366-specific effector CD8 T cells to the lung parenchyma, we performed intravascular staining with fluorophore-labeled CD45.2 antibodies (43); only vascular but not parenchymal lymphocytes are expected to bind to intravenously injected anti-CD45.2 antibodies. NP366-specific CD8 T cells were detected in both the lung vasculature (CD45.2^+ve^) and lung parenchyma (CD45.2^-ve^) in WT mice, but the vast majority (∼96%) of NP366-specific CD8 T cells localized to the lung parenchyma **(Figure 2B)**. As in WT mice, the majority of NP366-specific CD8 T cells were found in lung parenchyma of vaccinated CCR2^−/−^ mice, suggesting that CCR2 plays a dispensable role in regulating the vascular versus parenchymal localization of effector CD8 T cells in lungs.

We assessed whether CCR2 deficiency affected mucosal imprinting of effector CD8 T cells by examining expression of CD103 and CD69 by vaccine-induced NP366-specific CD8 T cells in lungs. While the expression of CD69 on NP366-specific CD8 T cells in airways and lungs was comparable in vaccinated WT and CCR2^−/−^ mice, percentages of CD103^+ve^ NP366-specific CD8 T cells in CCR2^-/-^ mice were significantly higher than in lungs of WT mice. These data suggested that monocytes might limit CD103 expression on vaccine-elicited effector CD8 T cells. **(Figure 2C-D).** Expression of CD103 and differentiation of T_RM_s are regulated by antigen receptor signaling and expression of transcription factors such as T-bet and TCF-1 (44–46). First, we quantified levels of transcription factors T-bet and TCF-1 in NP366-specific effector CD8 T cells in lungs of vaccinated WT and CCR2^-/-^ mice. The percentages of T-bet^+ve^ NP366-specific effector CD8 T cells were significantly lower in lungs of CCR2-/- mice, as compared to those in WT mice (**Figure 2E-F**). Further, the expressions levels of T-bet but not TCF-1 (measured by median fluorescence intensities [MFI]) in CCR2^-/-^ NP366-specific CD8 T cells were significantly lower (*P*<0.05) than in their WT counterparts; TCF-1:T-bet ratios in CCR2-/- CD8 T cells were significantly higher than in WT CD8 T cells (**Supplementary Figure 2**). These data suggested that CCR2-dependent pulmonary monocyte infiltration limits mucosal imprinting of effector CD8 T cells by inducing T-bet expression.

ADJ is known to drive strong T cell receptor (TCR) signaling and terminal differentiation of effector cells, while adding GLA to ADJ dampens TCR signaling and terminal differentiation of effector cells in the respiratory tract (40). Since PD-1 expression can serve as a qualitative readout for TCR signaling in lungs of influenza-infected mice (47), we compared PD-1 expression on NP366-specific CD8 T cells in lungs of vaccinated WT and CCR2^-/-^ mice. PD-1 expression by NP366-specific effector CD8 T cells in lungs and BAL was comparable in WT and CCR2^-/-^ mice (**Figure 2G-H**). To directly determine whether CCR2 deficiency affected antigenic stimulation of CD8 T cells in DLNs and lungs, we adoptively transferred naïve OVA SIINFEKL-specific TCR transgenic OT-I CD8 T cells that express eGFP under the control of Nur77 promoter; eGFP expression induced by the Nur77 promoter faithfully reports ongoing TCR signaling (48). Subsequently, mice were vaccinated with OVA formulated in ADJ+GLA, and eGFP expression by donor OT-I CD8 T cells in lungs and DLNs was quantified by flow cytometry. Here, we found that Nur77-eGFP expression by OT-I CD8 T cells in DLN and/or lungs of WT and CCR2^-/-^ mice was comparable on days 2, 5, and 8 after vaccination. These data suggested that CCR2 deficiency did not significantly affect antigenic stimulation of T cells in the respiratory tract **(Figure 2I).** Thus, enhanced expression of CD103 and reduced T-bet levels in CCR2^-/-^ effector CD8 T cells cannot be explained by altered TCR signaling or expressions of PD-1, at least in the first 8 days after vaccination.

To determine the differentiation state of vaccine-induced NP366-specific effector CD8 T cells in respiratory tract, we quantified CD127 and KLRG-1 expression and classified them as: short-lived effector cells (SLECs; CD127^LO^/KLRG-1^HI^), memory precursor effector cells (MPECs; CD127^HI^/KLRG-1^LO^), transition effector cells (TEs; CD127^Hi^/ KLRG1^HI^), and early effector cells (EEs; CD127^LO^/ KLRG1^Lo^). A substantive fraction of NP366-specific CD8 T cells were MPECs in airways and lungs of WT mice, but the relative proportions of CD127^Hi^/KLRG-1^lo^ MPECs were significantly (*P*<0.05) higher in CCR2^−/−^ mice, as compared to WT mice, suggesting that monocytes might restrain the development of MPECs in lung **(Figure 2J-K).** We further examined the differentiation status of effector CD8 T cells in CCR2^-/-^ mice by measuring expression of CD62L and OX40. In both airways and lungs, NP366-specific effector CD8 T cells in CCR2-/- mice exhibited significantly (*P*<0.05) increased expression of OX40 and CD62L **(Supplementary Figure 3)**, as compared to those in WT mice. Taken together, elevated expression of CD127, CD62L and OX40 in NP366-specific CD8 T cells in CCR2^-/-^ mice suggested that monocytes might promote the differentiation of KLRG-1^HI^/CD62L^LO^/OX40^LO^ effector CD8 T cells in the lungs.

### Effect of CCR2 deficiency on vaccine-induced effector CD4 T cells in the respiratory tract

Here, we asked whether CCR2 deficiency and loss of monocytes affected the accumulation and differentiation of vaccine-induced effector CD4 T cells in the respiratory tract. At day 8 after booster vaccination with ADJ+GLA+NP, high percentages of NP311-specific CD4 T cells were detected in airways and lungs of both WT and CCR2-/- mice **(Figure 3A).** The percentages and total numbers of NP311-specific CD4 T cells in lungs and airways were comparable between WT and CCR2^-/-^ mice. Furthermore, ∼96% of effector NP311-specific CD4 T cells were found in the lung parenchyma of both WT and CCR2^-/-^ mice **(Figure 3B).** Unlike for effector CD8 T cells (**Figure 2**), CCR2 deficiency did not affect mucosal imprinting of effector CD4 T cells in lungs; percentages of CD103^+ve^ cells amongst NP311-specific effector CD4 T cells were comparable in WT and CCR2^-/-^ mice **(Figure 3C-D)**. However, CCR2 deficiency promoted the development of KLRG-1^LO^/CD127^HI^ (MPECs) and CD62L^+ve^/OX40^+ve^ NP311-specific effector CD4 T cells in lungs and airways of vaccinated mice **(Figure 3E-H)**. Thus, CCR2 and monocytes might promote terminal differentiation of effector CD4 T cells in vaccinated mice.

**Figure 3.**
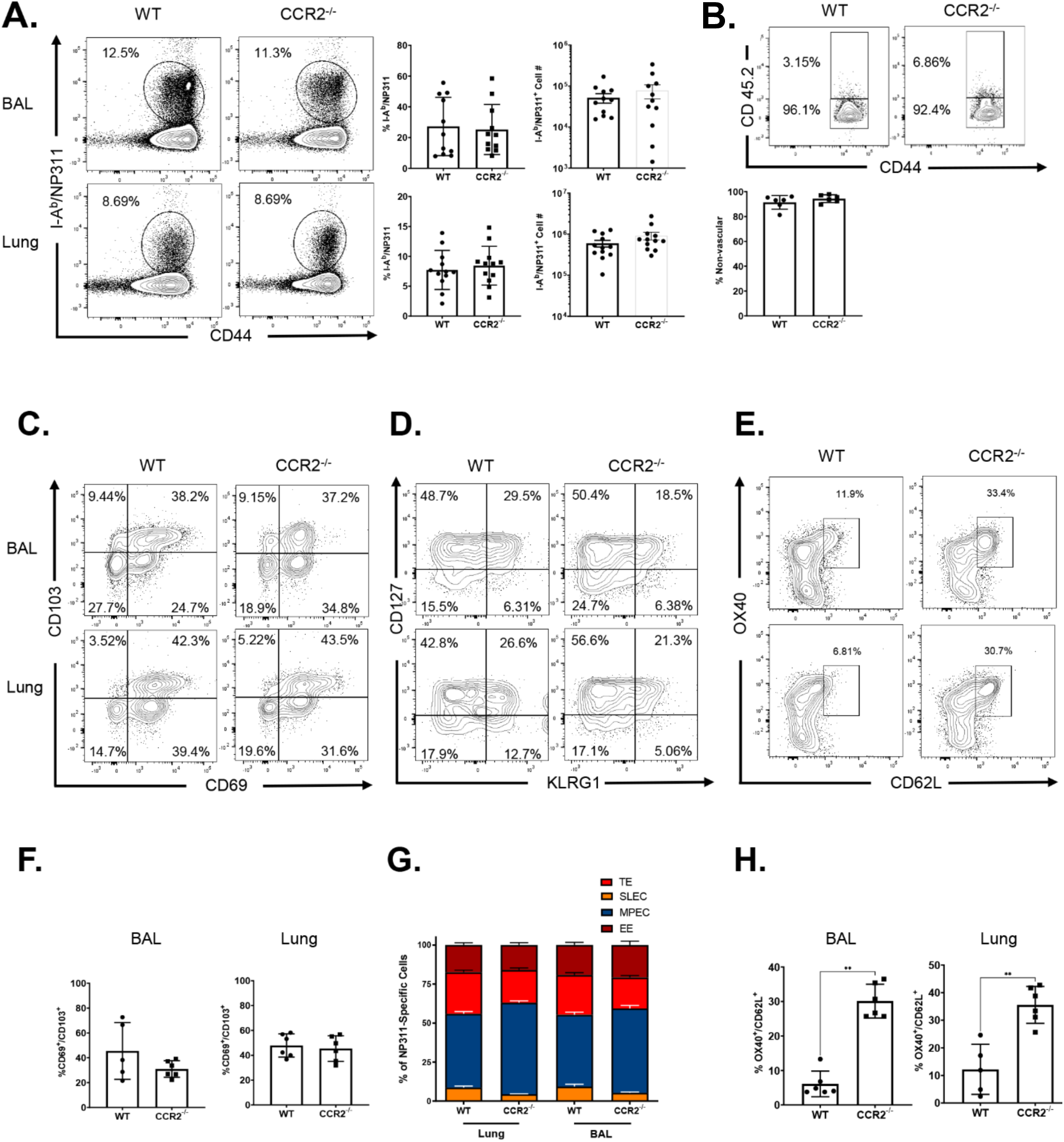
Effector CD4 T cell responses to adjuvanted subunit vaccine in CCR2^-/-^ mice. Wild Type (WT) or CCR2^-/-^mice were vaccinated as described in Figure 2. To distinguish non-vascular cells from vascular cells in the lungs, mice were injected intravenously with fluorescent-labeled anti-CD45.2 antibodies, 3 min prior to euthanasia (CD45.2^+ve^ – vascular; CD45.2^-ve^ – non-vascular. On the 8^th^ day after vaccination, single-cell suspensions prepared from the lungs and bronchoalveolar lavage (BAL) were stained with viability dye, followed by I-A^b^ /NP311 tetramers in combination with anti-CD4, anti-CD8, anti-CD44, anti-CD69, anti-KLRG1, anti-CD127, anti-OX40, and anti-CD62L antibodies (A, C-H). (A) FACS plots show percentages of tetramer-binding cells among CD4 T cells. (B) Percentages of vascular and non-vascular cells among NP311-specific CD4 T cells. Data are pooled from two independent experiments (A) or represent one of two independent experiments (B-H). Mann-Whitney U test, *, **, and *** indicate significance at *P*<0.1, 0.01 and 0.001 respectively.

### Functional polarization of vaccine-induced mucosal effector CD8 and CD4 T cells in CCR2-/- mice

As an index of effector differentiation, we measured granzyme B levels in NP366-specific effector CD8 T cells in lungs of vaccinated WT and CCR2^-/-^ mice, at day 8 after booster vaccination. Expression of granzyme B in NP366-specific CD8 T cells was not significantly different (*P*<0.05) in lungs of vaccinated WT and CCR2^-/-^ mice **(Figure 4A).** We have previously reported that vaccination with ADJ+GLA+NP fostered a functionally multifaceted T_C_1/T_C_17/T_H1_/T_H_17 response in lungs (40). To investigate the role of lung monocyte recruitment in the polarization of T_C_1/T_C_17 effector CD8 T cells, we vaccinated cohorts of WT and CCR2-/- mice twice, and assessed ex vivo cytokine production by NP366-specific CD8 T cells, at day 8 after booster vaccination. Upon *ex vivo* NP366 peptide stimulation, NP366-specific effector CD8 T cells from lungs of WT and CCR2^-/-^ mice produced IFNγ and/or IL-17α **(Figure 4B).** CCR2 deficiency did not affect the percentages of IFNγ and/or IL-17α-producing CD8 T cells in lungs of vaccinated mice (**Figure 4B**). Likewise, CCR2 deficiency did not affect the polyfunctionality of NP366-specific effector CD8 T cells as measured by their ability to co-produce IFNγ, IL-2 and TNFα **(Figure 4C).** Furthermore, antigen-triggered production of GM-CSF by NP366-specific effector CD8 T cells in WT and CCR2-/- mice was similar **(Figure 4D)**. Similar to NP366-specific effector CD8 T cells, CCR2 deficiency did not alter the ability of NP311-specific CD4 T cells to produce IFNγ, IL-17α, TNFα, IL-2 or GM-CSF (**Figure 5**). In summary, functional polarization of T_C_1/T_C_17/T_H_1/T_H_17 was not affected by lack of CCR2 or monocyte recruitment into lungs of vaccinated mice.

**Figure 4.**
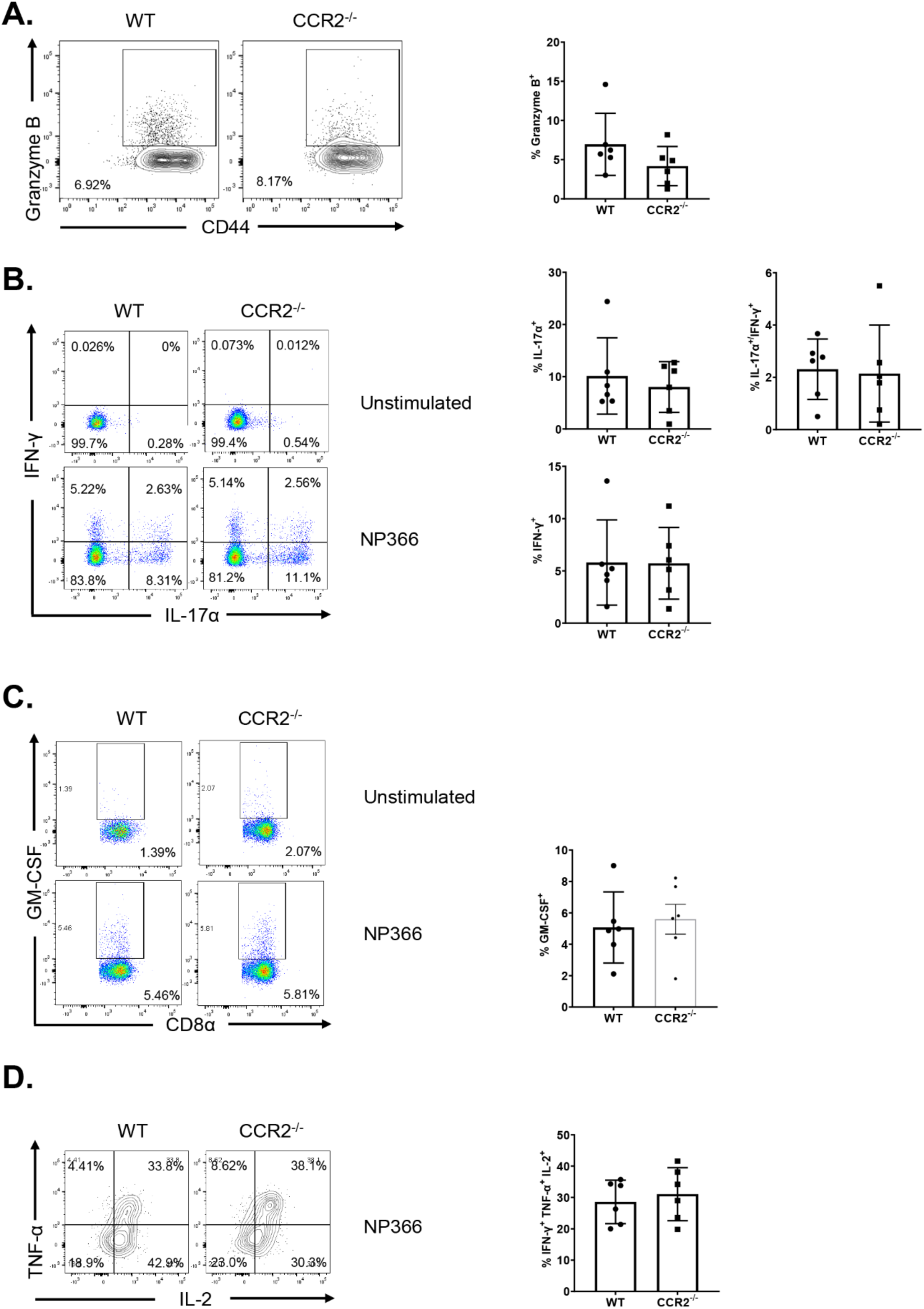
Functional polarization of effector CD8 T cells in vaccinated CCR2^-/-^mice. Wild Type (WT) or CCR2^-/-^mice were vaccinated as described in Figure 2. On the 8^th^ day after vaccination, lung cells were cultured with NP366 peptide, recombinant IL-2 and Brefeldin A for 5 h. The percentages of NP366-specific CD8 T cells that produced IL-17-α, IFN-γ, IL-2, and TNF-α, IL-10 and GM-CSF were quantified by intracellular cytokine staining. (A) Percentages of IFN-γ/IL-17α-producing cells among the gated CD8 T cells. (B) Percentages of IL-2/TNF-αproducing cells among the gated IFN-γ-producing CD8 T cells. (C) Percentages of GM-CSF-producing cells among the gated CD8 T cells. Cultures without the NP366 peptide (Unstimulated) served as a negative control. Data in each graph indicate Mean ± SEM. Mann-Whitney U test, *, **, and *** indicate significance at *P*<0.1, 0.01 and 0.001 respectively. Each independent experiment had n=3-5 mice per group. Data represent one of two independent experiments.

**Figure 5.**
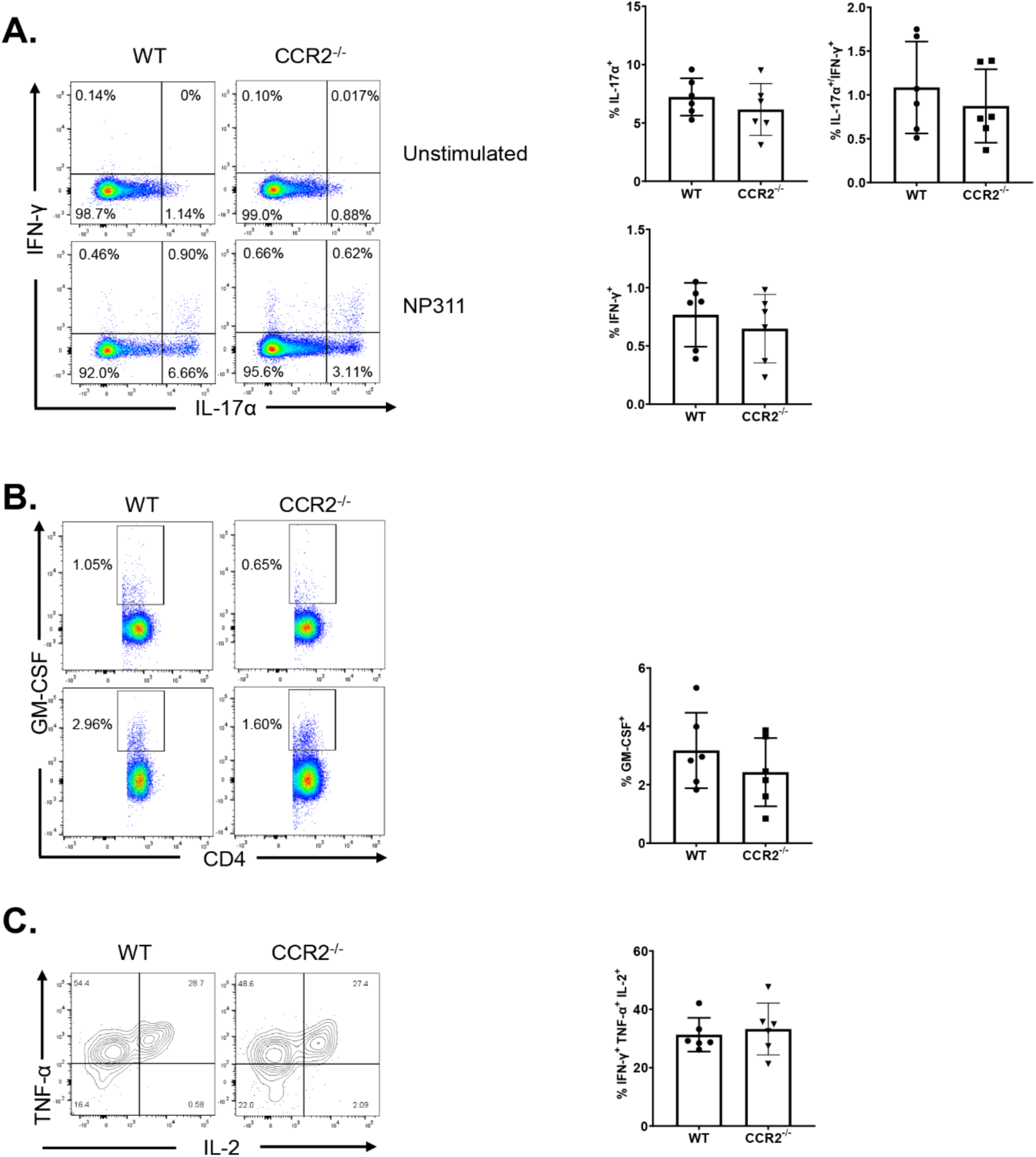
Functional polarization of effector CD4 T cells in vaccinated CCR2^-/-^mice. Wild Type (WT) or CCR2^-/-^mice were vaccinated, as described in Figure 2. On the 8^th^ day after vaccination, lung cells were cultured with NP311 peptide, recombinant IL-2 and Brefeldin A for 5 h. The percentages of NP311-specific CD4 T cells that produced IL-17-α, IFN-γ, IL-2, and TNF-α, IL-10, and GM-CSF were quantified by intracellular cytokine staining. (A) Percentages of IFN-γ/IL-17α-producing cells among the gated CD4 T cells. (B) Percentages of IL-2/TNF-α producing cells among the gated IFN-γ-producing CD4 T cells. (C) Percentages of GM-CSF-producing cells among the gated CD4 T cells. Cultures without the NP311 peptide (Unstimulated) served as a negative control.. Data in each graph indicate Mean ± SEM. Mann-Whitney U test, *, **, and *** indicate significance at P<0.1, 0.01 and 0.001 respectively. Each independent experiment had n=3-5 mice per group. Data represent one of two independent experiments.

### Mucosal CD8 and CD4 T cell memory in CCR2^-/-^ mice

Data in Figure 2 and 3 demonstrated that CCR2 deficiency augmented the expression of CD103 on NP366-specific effector CD8 T cells, but not NP311-specific effector CD4 T cells. Therefore, we assessed whether alteration of mucosal imprinting of effector CD8 T cells affected the development of T_RM_s in lungs and airways. At 60 days post-vaccination, the frequencies and numbers of NP366-specific memory CD8 T cells or NP311-specific memory CD4 T cells in lungs and airways of WT and CCR2^-/-^ mice were largely comparable **(Figure 6A, 6C).** Notably, the percentages of CD103^+ve^CD69^+ve^ NP366-specific memory CD8 T cells in lungs and airways of CCR2^-/-^ mice were significantly (*P*<0.05) higher than in WT mice. However, increased levels of CD103 on NP366-specific memory CD8 T cells in CCR2^-/-^ mice minimally affected their localization in the lung parenchyma of vaccinated mice (**Figure 6B).** In contrast to NP366-specific memory CD8 T cells, CCR2 deficiency neither affected CD103 expression or parenchymal localization of NP311-specific memory CD4 T cells in lungs or airways of vaccinated mice **(Figure 6E-F).** Thus, increased expression of CD103 induced by CCR2 deficiency in effector CD8 T cells was sustained in T_RM_s in lungs and airways of vaccinated mice.

**Figure 6.**
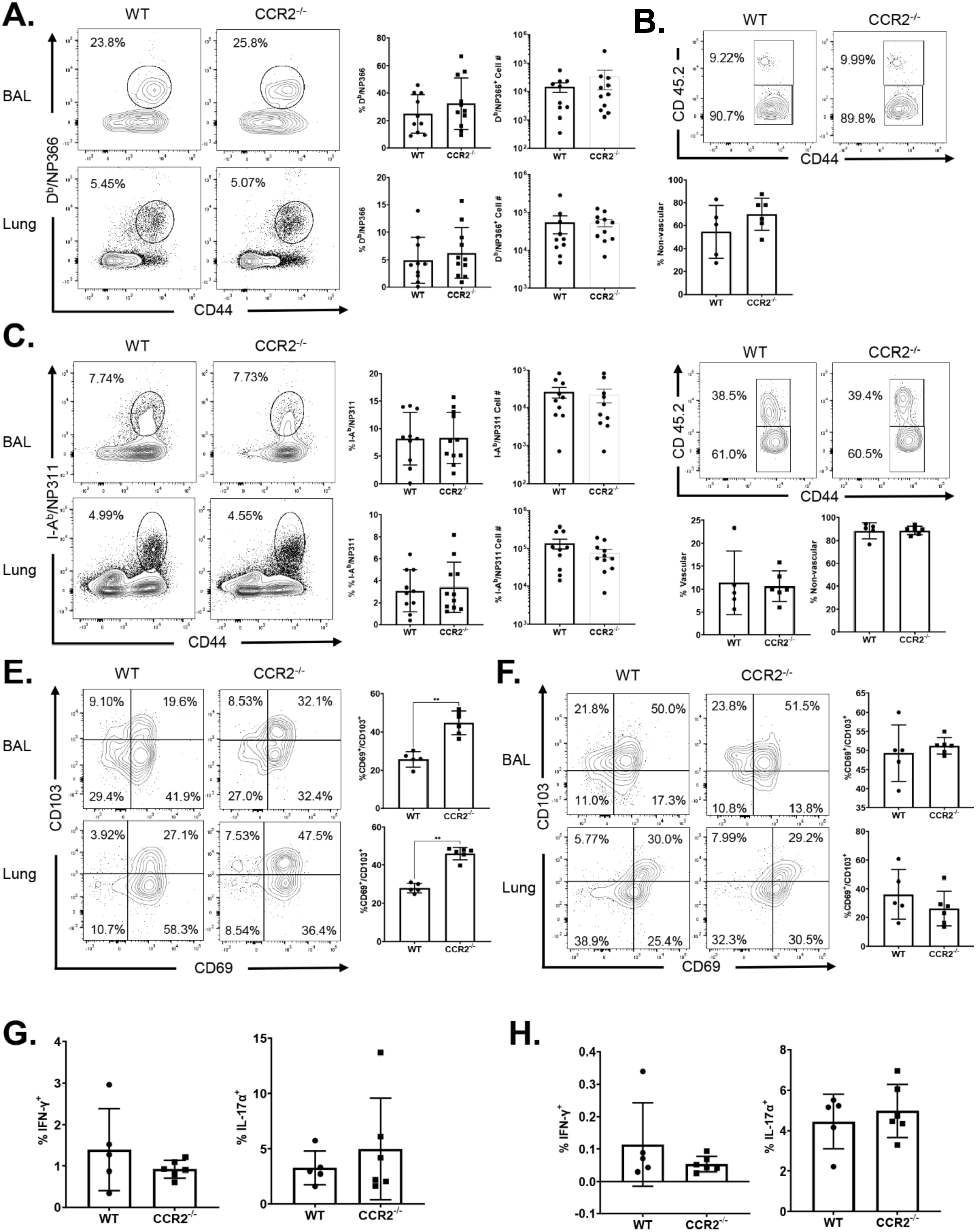
Vaccine-induced CD8 and CD4 T Cell Memory in CCR2^-/-^ mice. Wild Type (WT) and CCR2^-/-^ mice were vaccinated, as described in Figure 2. At 60 days after booster vaccination, the percentages and total numbers of NP366-specific memory CD8 T cells (A) and NP311-specific CD4 T cells (C) were quantified in lungs and airways (BAL). To distinguish non-vascular cells from vascular cells in the lungs, mice were injected intravenously with fluorescent-labeled anti-CD45.2 antibodies, 3 min prior to euthanasia (CD45.2^+ve^ – vascular; CD45.2^-ve^ – non-vascular). Single-cell suspensions from lungs or BAL were stained with D^b^/NP366, I-A^b^ /NP311, anti-CD8, anti-CD44, anti-CD103 and anti-CD69. All FACS plots in this figure are gated on tetramer-binding CD8 or CD4 T cells. (A or C) Percentages and total numbers of NP366-specific CD8 (A) or NP311-specific CD4 (B) T cells in BAL and Lung. (B or D) Percentages of vascular and non-vascular cells among NP366-specific CD8 (B) or NP311-specific CD4 (D) T cells. (E,F) CD69^+^ and/or CD103^+^ cells among NP366-specific CD8 (E) or NP311-specific CD4 (F) T cells in lungs. (G, H) Lung cells were cultured with or without NP366 (G) or NP311 peptides in the presence of recombinant IL-2 and Brefeldin A for 5 h. The percentages of NP366-specific CD8 T cells or NP311-specific CD4 T cells that produced IL-17αand/or IFN-γ were quantified by intracellular cytokine staining. Data in each graph indicate Mean ± SEM. Mann-Whitney U test, *, **, and *** indicate significance at P<0.1, 0.01 and 0.001 respectively. Each independent experiment had n=3-6 mice per group. Data are pooled from two independent experiments (A, C) or represent one of two independent experiments (B, D, E-H).

At 60 days after vaccination, we investigated whether functional polarization into T1/T17 effectors was maintained in memory T cells from CCR2^-/-^ mice. Upon ex vivo antigenic stimulation, NP366-specific memory CD8 T cells and NP311-specific memory CD4 T cells from lungs of WT and CCR2^-/-^ mice readily produced IFNγ and/or IL-17α. The percentages of memory T_C_1/T_C_17/T_H_1/T_H_17 cells were comparable in WT and CCR2^-/-^ mice **(Figure 6G-H)**. Hence, CCR2 deficiency did not affect the maintenance of T1/T17 programming in memory CD8 or CD4 T cells of vaccinated mice.

### Vaccine-induced pulmonary T-cell immunity to influenza A virus in CCR2^-/-^ mice

At 50-60 days after booster vaccination, we challenged vaccinated and unvaccinated WT and CCR2^-/-^ mice with a lethal dose of the mouse-adapted PR8/H1N1 IAV and quantified recall T cell responses and viral titers in the lungs at D6 post virus challenge. As expected, lungs of unvaccinated WT and CCR2^-/-^ mice contained high IAV titers (**Figure 7A**) and mice lost 10-15% of body weight after viral challenge (**Figure 7B**). The lungs of vaccinated WT and CCR2^-/-^ mice contained up to 6 logs lower viral burden, as compared to those in unvaccinated groups, and vaccinated mice did not exhibit detectable weight loss after viral challenge (**Figure 7A-B**). These data suggested that CCR2 function and monocyte recruitment are dispensable for vaccine-induced memory T cell-dependent control of IAV in mice. We then quantified recall CD8 and CD4 T cell responses in the lungs at day 6 after PR8/H1N1 challenge. The percentages and total number of recall NP366- and NP311-specific CD8 and CD4 T cells in lungs were comparable between WT and CCR2^-/-^ groups **(Figure 7C-D)**. Likewise, expression of the effector molecule granzyme B was strong but comparable in NP366-specific CD8 T cells from lungs of WT and CCR2^-/-^ mice (**Figure 8A**). We also compared antigen-induced cytokine-producing ability of NP-specific recall CD8 and CD4 T cells. NP366-specific CD8 T cells and NP311-specific CD4 T cells in WT and CCR2^-/-^ mice produced readily detectable but comparable levels of IL17-[box], IFN-γ, TNF-α, IL-2, and GM-CSF **(Figure 8B-D; Supplementary Figure 4)**. Taken together, data in Figure 7 and 8 indicated that CCR2 and monocyte recruitment are dispensable for vaccine-induced T-cell-dependent protective immunity to IAV.

**Figure 7.**
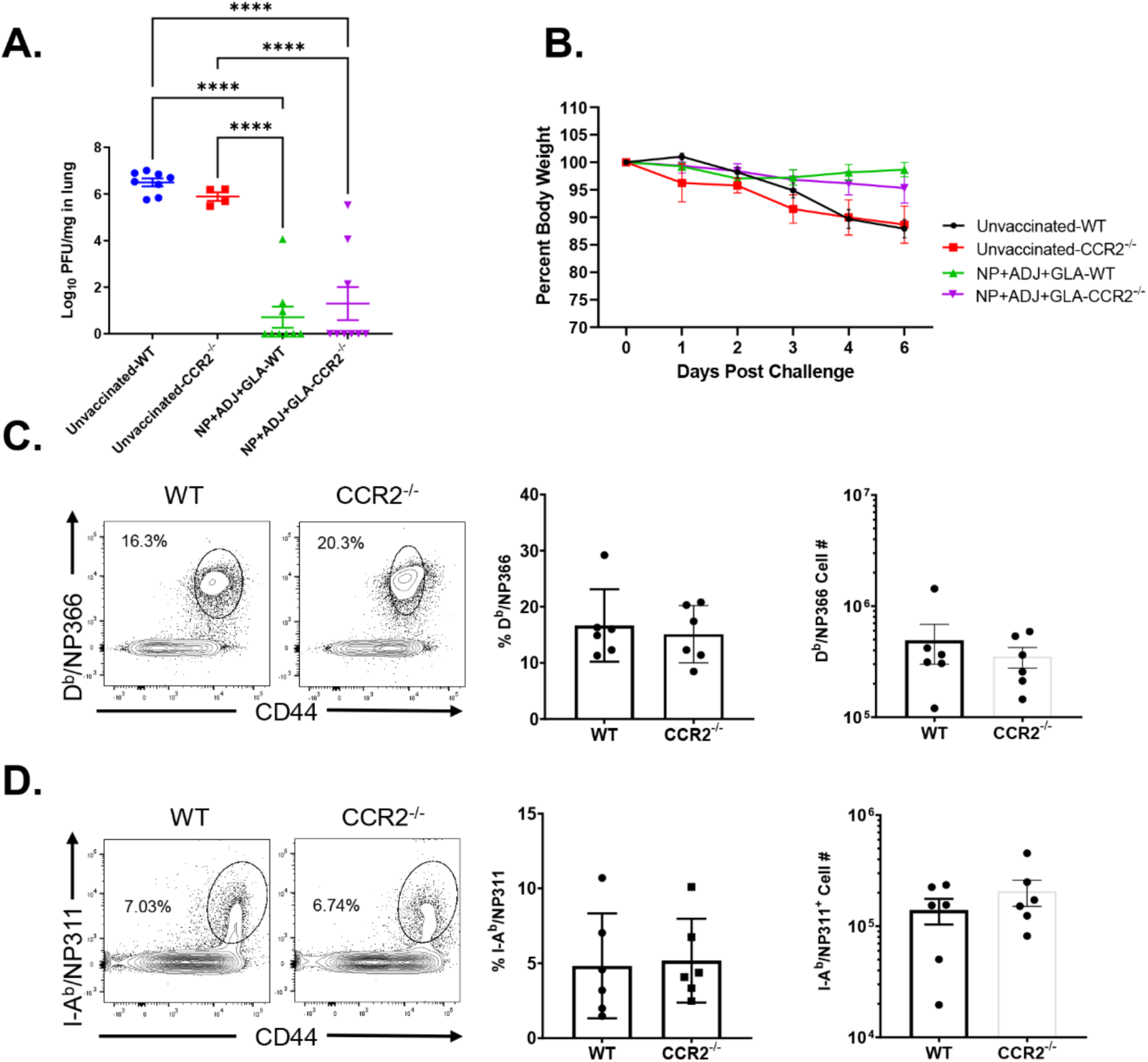
Vaccine-induced protective immunity to influenza A virus in WT and CCR2^-/-^ mice. At 50-60 days after booster vaccination, wild type (WT) or CCR2-/- mice were intranasally challenged with 500 PFUs of H1N1/PR8 strain of influenza A virus. (A) Viral titers were quantified in the lungs on D6 after challenge. (B) Following viral challenge, body weights were measured and plotted as a percentage (%) of starting body weight prior to challenge. Single-cell suspensions prepared from lungs and bronchoalveolar lavage (BAL) were stained with viability dye, followed by D^b^/NP366 and I-A^b^ /NP311 tetramers in combination with anti-CD4, anti-CD8 and anti-CD44. (C, D) Frequencies and total number of NP366-specific CD8 (C) or NP311-specific CD4 (D) T cells in lungs. FACS plots are gated on total CD8 (C) or CD4 (D) T cells. Each independent experiment had n=3-6 mice per group. Data in each graph indicate mean ± SEM. The data are pooled from two independent experiment (A-B) or represent one of two independent experiments (C-F). Mann-Whitney U test, *, **, and *** indicate significance at P<0.1, 0.01 and 0.001 respectively.

**Figure 8.**
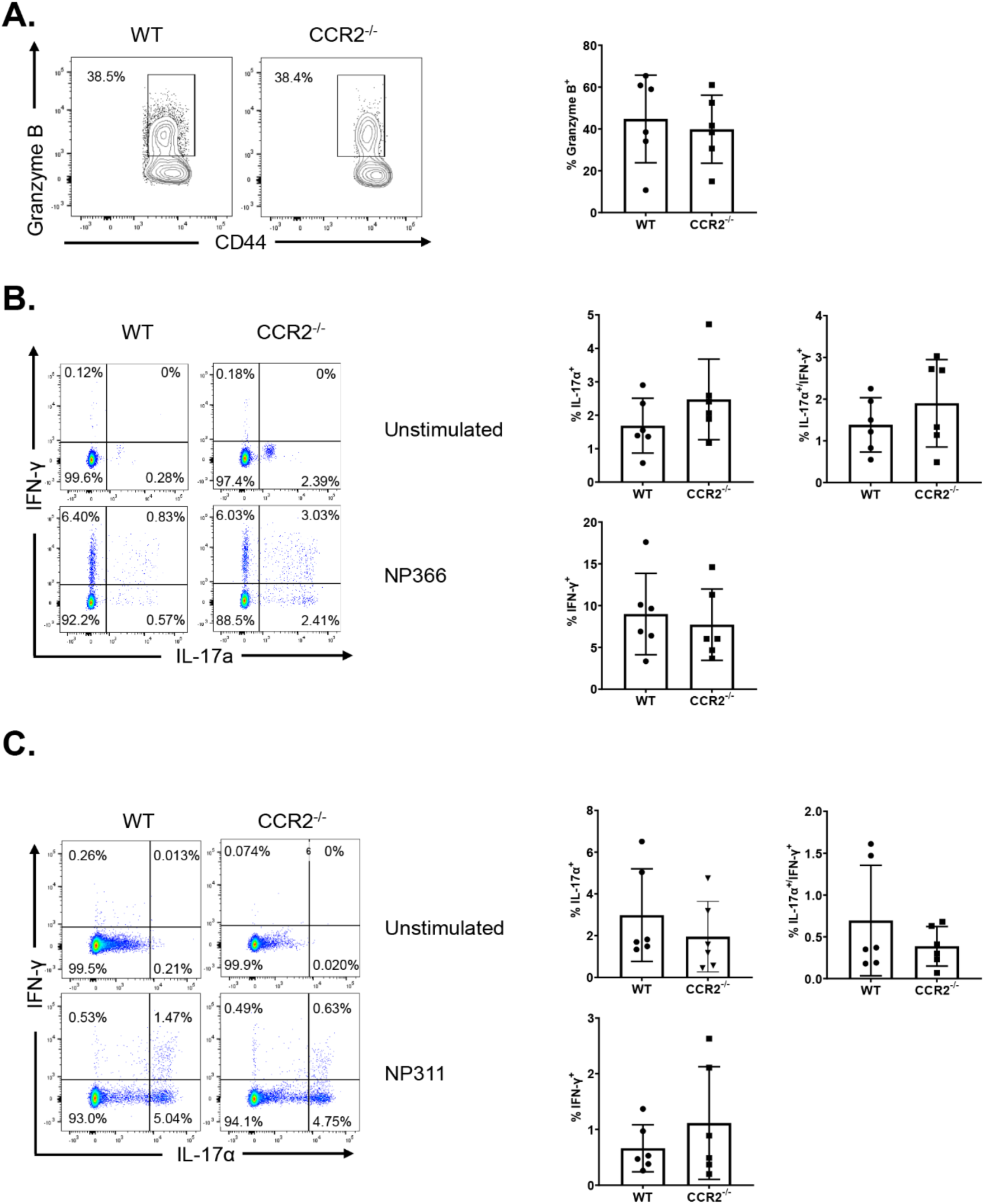
Functional characterization of recall CD4 and CD8 T cells in WT and CCR2^-/-^mice. At 50-60 days after booster vaccination, wild type (WT) or CCR2^-/-^ mice were challenged with H1N1/PR8 strain of influenza A virus. On the 6^th^ day after viral challenge, functions of antigen-specific CD4 and CD8 T cells in lungs were analyzed. (A) Single cell suspensions of lungs were stained directly ex vivo with D^b^/NP366 tetramers along with anti-CD8, anti-CD44 and anti-granzyme B antibodies. FACS plots in A are gated on tetramer-binding CD8 T cells and numbers are percentages of granzyme B^+ve^ cells among the gated population. (B, C) Single-cell suspensions of lungs were cultured with NP366 or NP311 peptides, and IFN-γ and/or IL-17α-producing CD8 or CD4 T cells were quantified by intracellular cytokine staining. Plots in B and C are gated on total CD8 and CD4 T cells, respectively. Numbers are percentages of IFN-γ and/or IL-17-α-producing cells among the gated population. Data are representative of two independent experiments.

## Discussion

Lung T_RM_s are a subset of memory T cells that reside in airways and lung parenchyma to provide first line of antigen-specific T cell defense against respiratory pathogens (10, 11, 21). Although it is well established that conventional DCs are crucial for initiating T cell priming (38, 49, 50), there are growing lines of evidence, suggesting a possible role for monocytes in influencing the differentiation and persistence of T_RM_s following recovery from respiratory viral infection (34, 35). In the current study, we have systematically documented the role of CCR2 and monocytes in orchestrating the differentiation of effector T cells, development of CD4 and CD8 T_RM_s, and recall responses following T-cell-based mucosal vaccination of mice. Data presented in this manuscript provide new insights into the role of innate immune cells, especially pulmonary monocytes in regulating mucosal imprinting and vaccine-induced T cell immunity in the respiratory tract.

Classic inflammatory monocytes are known to limit microbial invasion by secreting cytokines such as IL-1, IL-6, and TNF-α (51). Further, during a viral infection, under the influence of TLR agonists, inflammatory monocytes promote T_H_1 responses via direct priming of naïve T cells in the draining lymph node by cross-presentation (52–54). Importantly, antigen presentation by pulmonary monocytes to effector T cells appears to be vital for accumulation of effectors and development of T_RM_s in lungs of virally-infected mice (9, 34, 35). Further, IL-10-mediated TGF-β signaling induced by monocytes may have a critical role in the generation of T_RM_ following vaccination (55). Studies of IAV infection show that CCR2 is required for optimal accumulation of effector CD8 T cells in lungs and development of T_RM_s (9, 34, 35). Likewise, during a primary mucosal HSV-2 infection, activation of effector T cells in tissues was impaired in CCR2^-/-^ mice (56). In the HSV-1 reactivation model, inflammatory DCs (descendants of monocytes) were required to initiate memory responses in the tissue by way of activating CD4 and CD8 T_RM_s (57). However, we found that unlike an IAV or HSV-2 infection (9), CCR2 deficiency did not affect the magnitude of lung effector CD8 T cell responses to an adjuvanted mucosal vaccine or recall responses after viral challenge of vaccinated mice. Second, we found that CCR2 deficiency-induced loss of pulmonary monocytes led to enhancement of mucosal imprinting and development of CD103^+ve^CD69^+ve^ CD8 effector cells and T_RM_s in vaccinated mice. These findings suggest that mechanisms that regulate effector CD8 T cell accumulation and mucosal imprinting are likely different in virus infected versus vaccinated mice. Another inference is that the immunological milieu in the lungs of virus-infected and vaccinated mice is different, and that the environment in vaccinated mice can promote effector expansion and T_RM_ development in the absence of monocytes. Supporting this line of argument, it is noteworthy that IAV infection triggers a dominant T1-driving inflammation, but ADJ+GLA elicits an immunological milieu that fosters T1/T17 development. It is possible that effector expansion and development of T_RM_s in a T1 inflammatory environment but not in a T17-skewed milieu requires monocyte recruitment. Strikingly, the T1/T17-driving inflammatory milieu in lungs of ADJ+GLA vaccinated CCR2^-/-^ mice not only makes monocytes dispensable, it augmented mucosal imprinting and T_RM_ development in the absence of monocytes. Factors that govern mucosal imprinting or T_RM_ development include antigen, IL-10 and TGF-β (21, 55). We found that antigenic stimulation of CD8 T cells, as monitored by measuring Nur77-eGFP expression was not significantly different in DLNs and lungs of WT and CCR2^-/-^ mice. Therefore, it is less likely that enhanced mucosal imprinting in CCR2^-/-^ mice is driven by altered magnitude of TCR signaling in lungs. From the context of the inflammatory milieu, it is noteworthy that CCR2^-/-^ mice evoke a compensatory infiltration of neutrophils in the absence of monocyte recruitment (58). Because neutrophils are one of the major cell types that process vaccine antigen in lungs and can secrete active form of TGF-β1 (59, 60), it is plausible that neutrophils can act as another major cellular source of active TGF-β1 in CCR2^-/-^ mice. Follow-up studies should evaluate whether infiltration of neutrophils compensates for lack of monocytes and provide additional TGF-β signaling in CCR2^-/-^ mice, leading to augmented mucosal imprinting and development of lung T_RM_s.

It is also noteworthy that CCR2 deficiency was associated with increased levels of less differentiated CD127^Hi^KLRG-1^LO^ MPECs at the expense of the terminally differentiated CD127^LO^KLRG-1^HI^ SLECs. These findings suggest that CCR2 and pulmonary monocytes promote terminal differentiation of effector T cells in lungs of vaccinated mice. CD8 T cell terminal differentiation is typically driven by antigen receptor signaling and inflammation (45). It is possible that lack of pulmonary monocytes might have resulted in fewer antigen-presenting cells and reduced antigen encounter by T cells (9), but similar Nurr77-eGFP expression in OT-I CD8 T cells and PD-1 expression on effector CD8 T cells in WT and CCR2^-/-^ mice argue against this possibility. It is also plausible that loss of pulmonary monocytes in CCR2-/- mice results in impaired development of inflammatory monocyte-derived DCs (9), leading to dampened inflammatory milieu in the lungs. Since T_RM_s are believed to differentiate from CD127^HI^ cells (45), dampened inflammation and/or antigen receptor signaling in CCR2-/- mice might create an immunological milieu (with neutrophil-derived TGF-β) that prevents terminal differentiation but promotes development of T_RM_s. Inflammation drives T-bet expression and T-bet promotes terminal differentiation at the expense of T_RM_s (45, 46, 61). Therefore, decreased expression of T-bet in CCR2-/- effector CD8 T cells support the hypothesis that dampened pulmonary inflammation in the absence of monocytes/monocyte-derived DCs augments mucosal imprinting and T_RM_ development following vaccination. We have previously reported that ADJ is the vaccine component that promotes mucosal imprinting in lungs (40) and ADJ induces a metabolically quiescent state in cross-presenting DCs (38). It will be interesting to investigate whether migratory DCs in ADJ+GLA-vaccinated mice are less inflammatory and capable of augmented TGF-β-mediated preconditioning of CD8 T cells for T_RM_ fate in the draining lymph node (62). This mechanism might make monocytes dispensable or even a limiting factor for T_RM_ development in vaccinated mice.

The most effective vaccines to date protect by inducing high levels of neutralizing antibodies, but development of vaccines against diseases such as tuberculosis and AIDS has been a challenge for vaccinologists because immune defense against these diseases also require T cells. Hence, there is high interest in developing vaccine strategies that elicit robust and durable T cell immunity, especially in the mucosal tissues. We have previously reported that mucosal delivery of a subunit protein antigen formulated in a combination adjuvant (ADJ+GLA) elicited robust numbers of CD4 and CD8 T_RM_s in lungs and airways (40). The current study provided two unique insights into the mechanisms that regulate development of T_RM_s following mucosal administration of a vaccine formulated in this combination adjuvant. First, we show that mechanisms that regulate T_RM_ development are different for acute viral infections and vaccinations. Second, we uncover a negative regulatory role for pulmonary monocytes in driving mucosal imprinting and development of T_RM_s in lungs and airways, following mucosal vaccination. Results presented in this manuscript have improved our understanding of the mechanistic underpinnings of generating effective T cell-based protective immunity in the respiratory tract. These results might pave the way for the rational development of precision adjuvants to evoke frontline T cell immunity to respiratory pathogens at the mucosal frontiers.

---

**Supplementary Figure 1.**
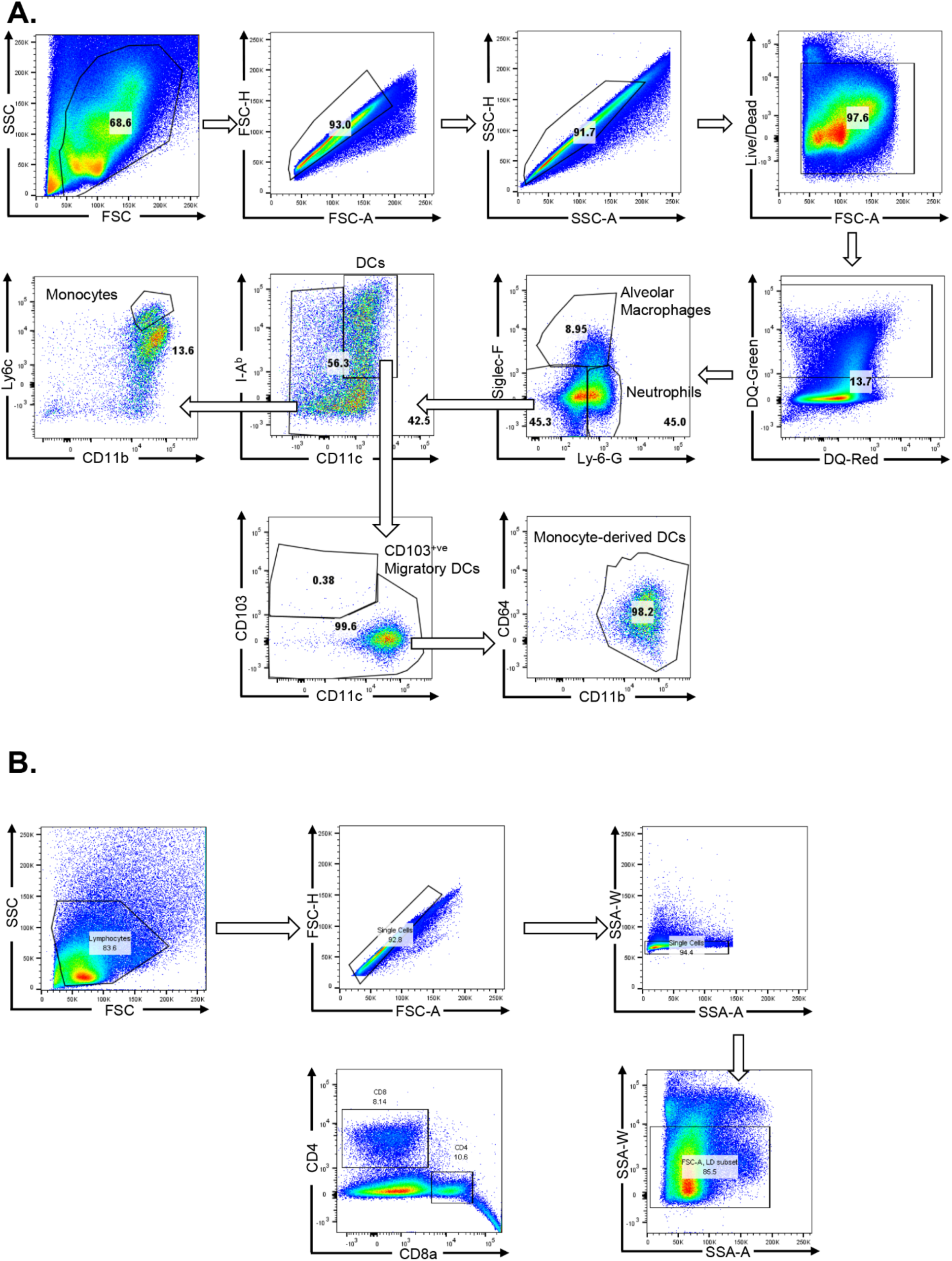
Analysis of innate immune cells and T cells in the lungs of vaccinated mice. (A) Gating strategy for innate immune cell subsets in lung. Groups of C57BL/6 mice were vaccinated with DQ-OVA protein (20ug) formulated in ADJ (5%) + GLA (5ug). At days 2, 5 and 8 after vaccination, lung cells were stained with anti-CD11b, anti-Siglec-F, anti-CD11c, anti-CD64, anti-Ly6G, anti-Ly6C, anti-CD103 and anti-I-A/I-E. Samples were immunophenotyped using the following parameters: neutrophils (Ly-6G^HI^/Siglec-F^LO^/CD64^LO^), alveolar macrophages Ly6G^LO^/Siglec-F^HI^/CD64^HI^CD103^LO^), monocytes (Ly6G^LO^/Siglec-F^LO^/MHC-II^Lo^/CD11c^LO^/CD64^LO^/CD103^LO^CD11b^H^/Ly6C^HI^), monocyte-derived DCs (Ly6G^LO^/SiglecF^LO^MHC-II/CD11c^HI^/CD64^HI^/CD103^LO^/CD11b^HI^/Ly6C^LO-INT^, and CD103^+ve^ migratory DCs (Ly6G^LO^/Siglec-F^LO^/CD64^LO^/MHC-II/CD11c^HI^/CD103^HI^/CD11b^LO^). (B) Gating strategy for visualization and analysis of antigen-specific T cells: FSC vs. SSC for lymphocyte gate → singlets → live-cell gate → CD4 or CD8 T cells.

**Supplementary Figure 2.**
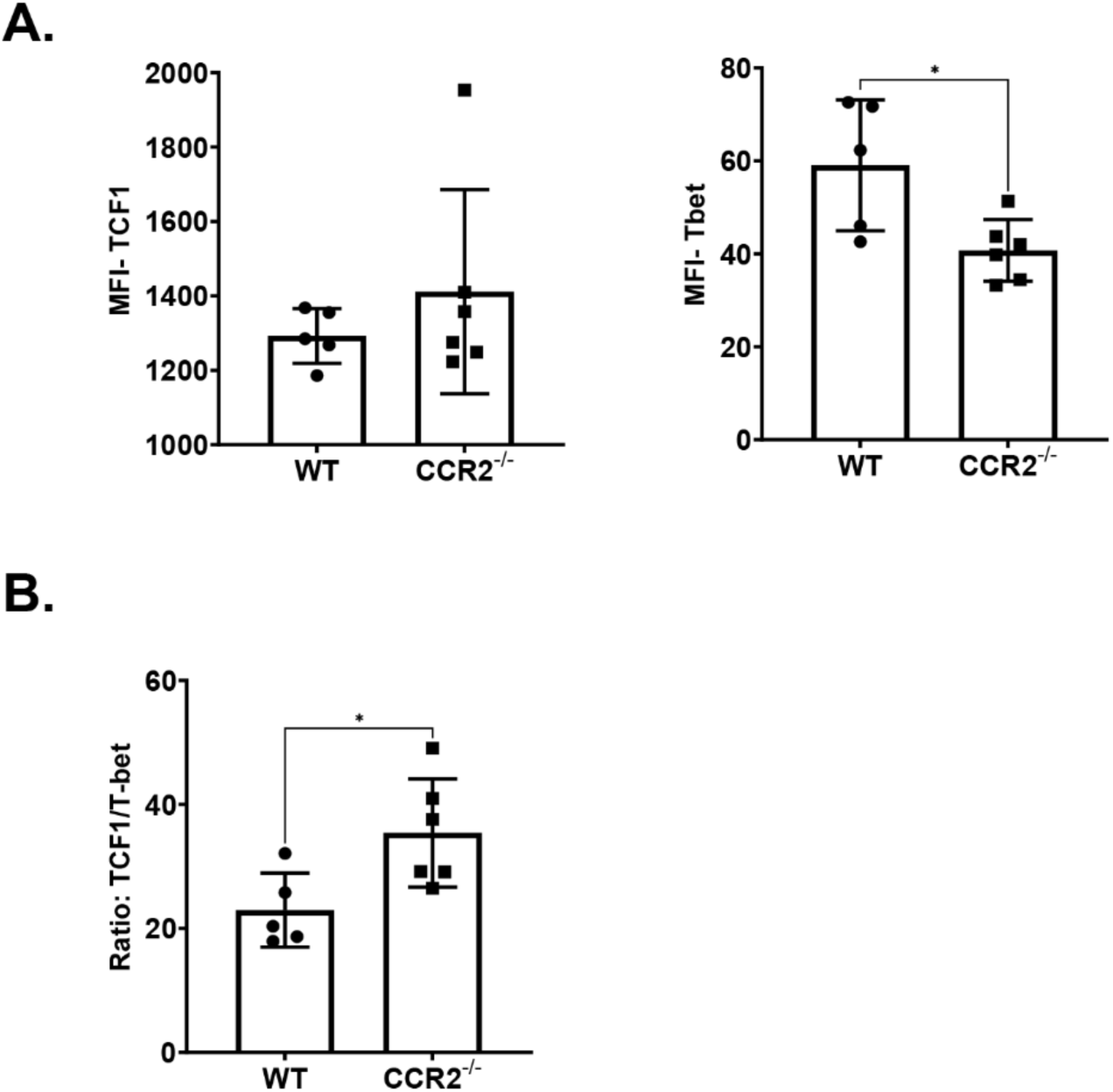
Expression of T-bet and TCF-1 in NP366-specific CD8 T cells. Wild type (WT) or CCR2^-/-^mice were vaccinated intranasally (IN) twice (21 day apart) with influenza A H1N1 Nucleoprotein (NP) formulated in ADJ (5%) and GLA (5ug). On the 8^th^ day after booster vaccination, single-cell suspensions from the lungs were stained with viability dye, followed by D^b^/NP366 tetramers in combination with anti-CD4, anti-CD8, and anti-CD44. The samples were subsequently permeabilized and stained with anti-TCF-1 and anti-T-bet. Samples were analyzed by flow cytometry. Data in A show median fluorescence intensities (MFI) for T-bet and TCF-1 in NP366-specific CD8 T cells. Panel B shows ratios of TCF-1:T-bet MFIs in each sample. Mann-Whitney U test, *, **, and *** indicate significance at P<0.1, 0.01 and 0.001 respectively.

**Supplementary Figure 3.**
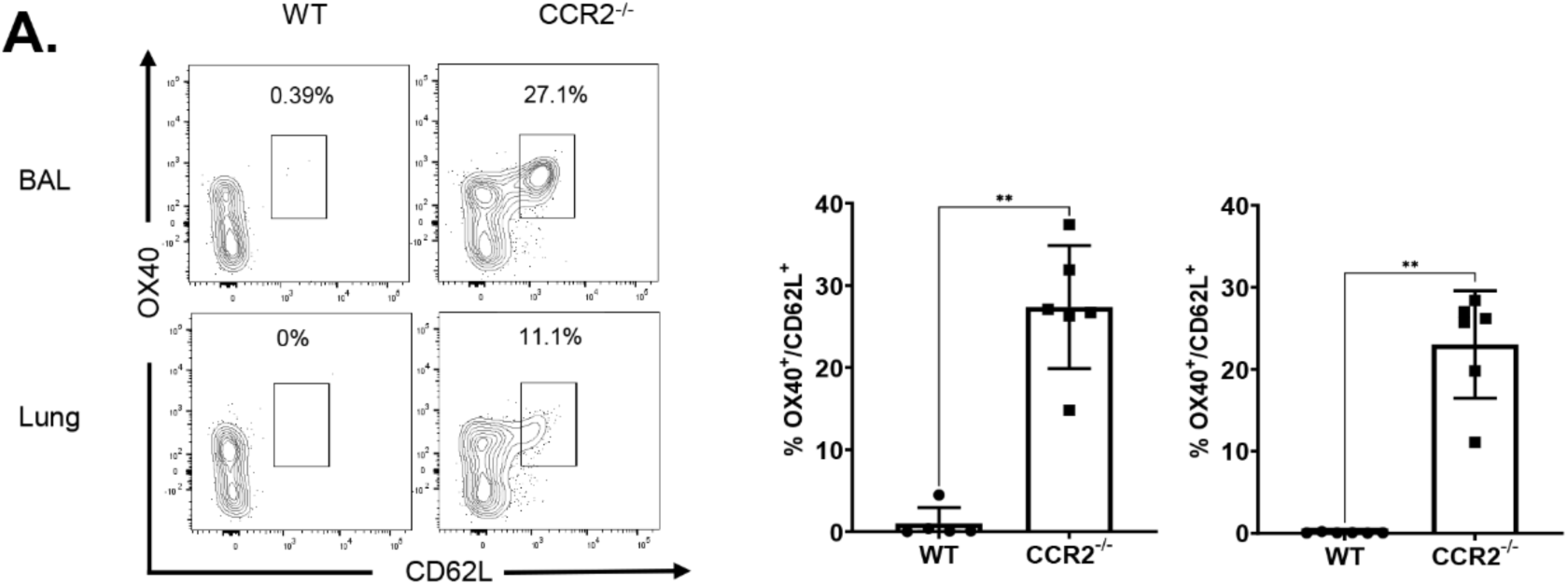
Expression of OX40 and CD62L in NP366-specific CD8 T cells. Wild type (WT) or CCR2^-/-^mice were vaccinated intranasally (IN) twice (21 day apart) with influenza A H1N1 Nucleoprotein (NP) formulated in ADJ (5%) and GLA (5ug). On the 8^th^ day after booster vaccination, single-cell suspensions from the lungs and bronchoalveolar lavage (BAL) were stained with viability dye, followed by D^b^/NP366 tetramers in combination with anti-CD4, anti-CD8, anti-CD44, anti-CD69, anti-CD103, anti-PD-1, anti-KLRG1, anti-CD127, anti-OX40 and anti-CD62L antibodies. FACS plots are gated on tetramer-binding CD8 T cells. Mann-Whitney U test, *, **, and *** indicate significance at P<0.1, 0.01 and 0.001 respectively.

**Supplementary Figure 4.**
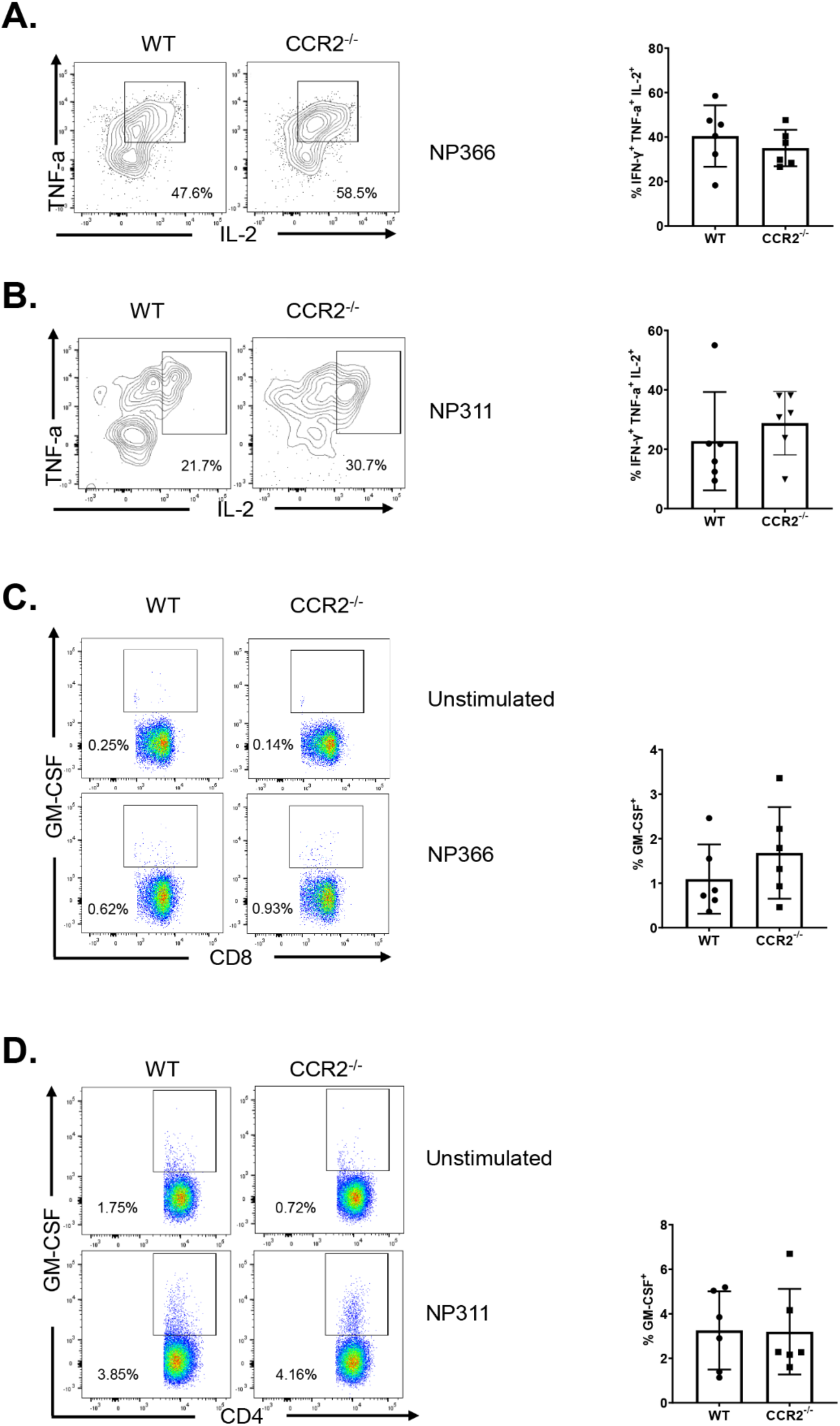
Functional polarization of recall CD8/4 T cells in WT and CCR2^-/-^mice. At 50-60 days after booster vaccination, WT or CCR2^-/-^ mice were challenged with H1N1/PR8 strain of influenza A virus. On the 6^th^ day after viral challenge, single-cell suspensions of lungs from challenged mice were *ex vivo* stimulated with NP366 or NP311 peptides. The percentages of NP366-stimulated CD8 T cells that produced IFN-γ, TNF-α, IL-2, and GM-CSF were quantified by intracellular cytokine staining. (A, B) IL-2- and TNF-α co-producing cells among the gated IFN-γ-producing CD4/8 T cells. (C, D) FACS plots are gated on total CD8 (C) or CD4 (D) CD4 T cells. Data are representative of two independent experiments. Mann-Whitney U test, *, **, and *** indicate significance at P<0.1, 0.01 and 0.001 respectively.

**Supplementary Table 1.** List of antibodies used in the manuscript.

| **Antibody (Dilution Factor)** | **Company** | **Catalogue number** |
| --- | --- | --- |
| Hamster anti-CD11c-BV786-conjugated (1:200) | BD Biosciences | 563735 |
| Rat anti-CD11b-BV711-conjugated (1:00) | BD Biosciences | 563168 |
| Rat anti- I-A/I-E-BV650-conjugated (1:400) | BD Biosciences | 563415 |
| Rat anti-Siglec-F-Alexa Fluor 647-conjugated (1:200) | BD Biosciences | 562680 |
| Rat anti-Ly-6G-BUV 395-conjugated (1:200) | BD Biosciences | 563978 |
| Rat anti-Ly-6C-PE-Cy7-conjugated (1:200) | BD Pharmingen | 560593 |
| Rat anti-CD8α-BUV395-conjugated (1:200) | BD Biosciences | 563786 |
| Rat anti-CD4-BUV496-conjugated (1:200), | BD Biosciences | 564667 |
| Rat anti-CD44-BV510-conjugated (1:200) | BD Biosciences | 563114 |
| Rat anti-CD62L-PE-CF594-conjugated (1:200) | BD Biosciences | 562404 |
| Hamster anti-CD69-PE-Cy7-conjugated (1:200) | BD Biosciences | 553237 |
| Hamster Anti-KLRG1-BV711-conjugated (1:200) | BD Biosciences | 564014 |
| Hamster Anti-CD279 (PD-1)-BV 650-conjugated (1:200) | BD Pharmingen | 744546 |
| Rat anti-IFN-γ-APC-conjugated (1:400) | BD Biosciences | 554413 |
| Rat anti- TNF-α-BV421-conjugated (1:400 or 1:500) | BD Biosciences | 554413 |
| Rat anti-CD127-BV605-conjugated (1:200) | Biolegend | 135041 |
| Hamster anti-CD103-BV605-conjugated (1:200) | Biolegend | 121433 |
| Mouse anti-CX3CR1-BV785-conjugated (1:200) | Biolegend | SA011F11 |
| Rat anti-GM-CSF PE-Cy7-conjugated (1:200) | Biolegend | 505412 |
| Rat anti-IL-17A-FITC conjugated (1:200) | Biolegend | 506908 |
| Rat anti-CD127-PerCP/Cyanine5.5-conjugated (1:150) | Biolegend | 135022 |
| Rat anti-IL-2 PE/Dazzle 594-conjugated (1:200) | Biolegend | 503840 |
| Mouse anti-CD64 (FcγRI)-PerCP-Cy5.5-conjugated | Biolegend | 139308 |
| Rabbit TCF1-Alexa Fluor 488-conjugated (1:200) | Cell Signaling Technology | 6444S |
| Mouse anti-CD45.2-violetFlour 450-conjugated (3ug/mouse) | Tonbo Biosciences | 75-0454-U100 |
| Rat anti-OX40 (CD134)- PerCP-Cyanine5.5 conjugated (1:200) | Tonbo Biosciences | 65-1341-U025 |
| Purified Anti-Mouse CD16 / CD32 (Fc Shield) (2.4G2, 1:100) | Tonbo Biosciences | 70-0161-U500 |
| APC-conjugated H2-Kb tetramers bearing the ovalbumin peptide SIINFEKL | NIH Tetramer Core Facility at Emory University |  |
| BV421-conjugated I-Ab tetramers bearing the NP peptide NP311 (QVYSLIRPNENPAHK) | NIH Tetramer Core Facility at Emory University |  |
| APC-conjugated-H2-Kb tetramers bearing the NP peptide NP366 (ASNENMDTM) | NIH Tetramer Core Facility at Emory University |  |

## METHODS

### Experimental animals

7-12-week-old C57BL/6J (B6) were purchased from restricted-access SPF mouse breeding colonies at the University of Wisconsin-Madison Breeding Core Facility or from Jackson Laboratory. CCR2^-/-^ (Stock number: 004999) and BATF3^-/-^ (Stock number: 013755) mice were purchased from Jackson Laboratory. B6. Nur77-GFP OT-1 mice were bred in the laboratory of Dr. Ross M. Kedl (University of Colorado, Denver).

### Ethics statement

All experiments were performed in accordance with the animal protocol (Protocol number V5308 or V5564) approved by the University of Wisconsin School of Veterinary Medicine Institutional Animal Care and Use Committee. The animal committee mandates that institutions and individuals using animals for research, teaching, and/or testing much acknowledge and accept both legal and ethical responsibility for the animals under their care, as specified in the Animal Welfare Act and associated Animal Welfare Regulations and Public Health Service (PHS) Policy.

### Vaccination

Adjuplex (ADJ) and Glucopyranosyl Lipid Adjuvant (GLA) were purchased from Empirion LLC (Columbus, OH) and Avanti Polar Lipids, Inc. (Alabaster, AL), respectively. All vaccinations were administered intranasally to anesthetized mice in 50 ul saline with 10 ug NP (Influenza A H1N1 Nucleoprotein / NP Protein, Sino Biological) or 20ug DQ-OVA protein (Thermo Fisher Scientific) alone or with the following adjuvants: ADJ (5%)+GLA (5ug)

### Adoptive transfer of Nur77-eGFP/OT-I CD8 T Cells

Single-cell suspensions of spleens and lymph nodes (LNs) from Nur77-eGFP OT-I (CD-45.1^+ve^) mice containing 10^3^ (vaccinated mice) or 1×10^6^ (for unvaccinated mice) of transgenic CD8+ T cells were injected intravenously into sex-matched congenic CD45.2 C57BL/6 mice. 24 hours later, mice were intranasally vaccinated with OVA formulated with ADJ+GLA. At days 2, 5, and 8 after vaccination, cells from lungs and mediastinal lymph nodes were stained with anti-CD8, anti-CD45.1, anti-CD44 and K^b^/SIINFEKL tetramers. Nur77-eGFP expression by live OT-I CD8 T cells (gated on CD8, CD45.1, Nur77-eGFP) was quantified directly *ex vivo* by flow cytometry.

### Tissue processing and flow cytometry

Lungs and draining lymph nodes were processed using standard collagenase-based digestion, as previously described (40). Briefly, lung tissue was minced and processed using the gentleMACS Dissociator (Miltenyi Biotech) in 5 ml of 1% RPMI media containing 2mg/ml collagenase B, as per manufacturer’s instructions. Samples were incubated for 30 minutes at 37C, re-homogenized and resuspended in media containing 1% fetal bovine serum (FBS). Subsequently, cells were spun down, resuspended in RPMI media containing 10% FBS and counted in a hemocytometer. 100ul (peak or recall) or 200ul (memory time) of single cell suspensions of cells (10^7^/ml) prepared from various tissues were stained for viability with Dye eFluor 780 (eBiosciences, San Diego, CA) or Ghost Dye 780 (Tonbo Biosciences) and incubated with fluorochrome-labeled antibodies and MHC I tetramers (see supplementary table 1 for dilution) at 4C for 1 hour. For staining with the I-A^b^/NP311 tetramer (1:150 dilution), cells were incubated with tetramer at 37C for 90 minutes, followed by staining with antibodies indicated in **Supplementary table 1** for cell surface molecules at 4C for 60 minutes. Following staining, cells were washed twice with FACS buffer (2% BSA in PBS). Stained cells were fixed with 2% paraformaldehyde for 20 minutes, then transferred to FACS buffer (2% BSA in PBS). Data from live single cells were analyzed with FlowJo software (TreeStar, Ashland, OR).

### Influenza virus challenge studies and viral titration

For viral challenge studies, vaccinated mice were intranasally challenged with 500 plaque forming units (PFUs) of A/PR8/8/1934 (H1N1) strains of influenza A virus diluted in 50 uL PBS (39, 40). On the 6^th^ day after influenza challenge, lungs were collected from mice on the 6th day for viral titration by a plaque assay using Madin Derby Canine Kidney Cells (MDCK) cells as previously described (40).

### Intracellular staining for Granzyme B and transcription factors

To stain for granzyme B or transcription factors, single-cell suspension were first stained for viability with LiveDead eFlour 780 stain (eBioscience) or Ghost Dye™ Red 780 (Tonbo Biosciences) for 30 minutes and then stained with antibodies and tetramers diluted in Brilliant Stain Buffer (BD Biosciences) for 60 minutes. The samples were then fixed, permeabilized and subsequently stained for transcription factors using the transcription factors staining kit (eBioscience) with the antibodies indicated in **Supplementary table 1** in Perm Wash buffer. All samples were acquired on LSRFortessa (BD Biosciences) and analyzed with FlowJo V.10 software (TreeStar, Ashland, OR).

### Intracellular cytokine staining

For intracellular cytokine staining, one million cells (1×10^6^) cells were plated on flat-bottom tissue-culture-treated 96-well plates (Corning.) Cells were stimulated for 5 hours at 37C in the presence of brefeldin A (1 μl/ml, GolgiPlug, BD Biosciences), human recombinant IL-2 (10 U/well) and with or without NP311 or NP366 peptides (Genscript) at 0.2ug/ml. After *ex vivo* peptide stimulation, cells were stained for viability dye LiveDead eFlour 780 stain (eBioscience) or Ghost Dye™ Red 780 (Tonbo Biosciences) for 30 minutes, stained with surface antibodies, and fixed/permeabilizated with Cytofix/Cytoperm kit (BD Biosciences, Franklin Lakes, NJ) according to manufacturer’s protocol. Samples were stained with antibodies indicated in **Supplementary table 1** in perm wash buffer for 30 minutes, washed with perm wash buffer, and re-suspended in FACS buffer before flow cytometry.

### Statistical analyses

Statistical analyses were performed using GraphPad software 9 (La Jolla, CA). All comparisons were made using either Mann-Whitney U test or an one-way ANOVA test with Tukey corrected multiple comparisons where p<0.05 = *, p<0.005 = **, p<0.0005 = *** were considered significantly different among groups.

## Acknowledgements

We thank Autumn Larsen and Daisy Gates for expert technical assistance. We also are thankful to Dr. Chandranaik B. Marinaik for assistance with vaccination. We gratefully acknowledge Emory NIH Tetramer Core Facility for providing MHC-I and MHC-II tetramers. Many thanks to Dr. Lisa Arendt for providing CCR2^-/-^ strains to initiate this project. We would also like to thank genuine appreciation for the efforts of the veterinary and animal care staff at UW-Madison.

## Funding

This work was supported by PHS grant U01 AI124299, R21 AI149793-01A1 and John E. Butler professorship to M. Suresh. Woojong Lee was supported by a predoctoral fellowship from the American Heart Association (18PRE34080150).

## Author contributions

W.L, B.B and M.S. designed, performed, analyzed experiments, and provided conceptual input for the manuscript. Y.K and R.K provided critical reagents for the manuscript. W.L and M.S wrote the manuscript, which was proofread by all authors.

